# High fat diet allows food-predictive stimuli to energize action performance in the absence of hunger, without distorting insulin signaling on accumbal cholinergic interneurons

**DOI:** 10.1101/2023.03.29.534845

**Authors:** Joanne M. Gladding, Nura W. Lingawi, Beatrice Leung, Michael D. Kendig, Billy C. Chieng, Vincent Laurent

## Abstract

Obesity can disrupt how food-predictive stimuli control action performance and selection. These two forms of control recruit cholinergic interneurons (CIN) located in the nucleus accumbens core (NAcC) and shell (NAcS), respectively. Given that obesity is associated with insulin resistance in this region, we examined whether interfering with CIN insulin signaling disrupts how food-predictive stimuli control actions. To interfere with insulin signaling we used a high-fat diet (HFD) or genetic excision of insulin receptor (InsR) from cholinergic cells. HFD left intact the capacity of food-predictive stimuli to energize performance of an action earning food when mice were tested hungry. However, it allowed this energizing effect to persist when the mice were tested sated. This persistence was linked to NAcC CIN activity but was not associated with distorted CIN insulin signaling. Accordingly, InsR excision had no effect on how food-predicting stimuli control action performance. Next, we found that neither HFD nor InsR excision altered the capacity of food-predictive stimuli to guide action selection. Yet, this capacity was associated with changes in NAcS CIN activity. These results indicate that insulin signaling on accumbal CIN does not modulate how food-predictive stimuli control action performance and selection. However, they show that HFD allows food-predictive stimuli to energize performance of an action earning food in the absence of hunger.

## 1. Introduction

Brain insulin resistance is associated with reduced sensitivity and efficacy of the insulin receptor (InsR) (Arnold et al., 2014; Clegg et al., 2011), and is a key feature of metabolic disorders such as obesity (Kullmann et al., 2016). As they age, people with obesity are at a higher risk of cognitive decline and dementia (Bischof & Park, 2015). Accordingly, it is thought that brain insulin resistance can contribute to the metabolic and cognitive deficits observed in obesity (Cholerton, Baker, & Craft, 2013). Yet, the precise nature of these deficits and their underlying mechanisms are poorly understood.

People with obesity show altered insulin signaling in the nucleus accumbens (NAc) (Kullmann et al., 2016; Heni et al., 2017), a brain region that plays a critical role in metabolic and cognitive functions (Castro & Bruchas, 2019; Ferrario, 2020; Morales & Berridge, 2020; Floresco, 2015). In the laboratory, these functions can be studied through the general and specific form of the Pavlovian-instrumental transfer (PIT) task (Holmes, Marchand, & Coutureau, 2010; Cartoni, Balleine, & Baldassarre, 2016). General PIT demonstrates that a stimulus predicting a food outcome energizes performance of an action earning another food outcome. By contrast, specific PIT reveals that a stimulus predicting a particular food outcome guides choice towards an action earning the same, but not a different outcome. Evidence shows that metabolic needs control general PIT but not specific PIT. This is typically shown by manipulating the primary motivational state of animals undergoing general or specific PIT tests (Corbit, Janak, & Balleine, 2007; Lingawi, Berman, Bounds, & Laurent, 2022; Balleine, 1994; Rescorla, 1994; Sommer, Münster, Fehrentz, & Hauber, 2022). These manipulations show that general PIT is present in hungry animals but is absent in sated animals. By contrast, animals display specific PIT whether they are tested hungry or sated. This dissociation suggests that general PIT models the metabolic regulation of action performance whereas specific PIT studies a cognitive, rather than metabolic, control of action selection.

Importantly, in the context obesity and potential deficits in NAc functioning, the core territory of the NAc (NAcC) mediates general PIT whereas its shell territory (NAcS) is required for specific PIT (Corbit, Fischbach, & Janak, 2016; Corbit, Muir, Balleine, & Balleine, 2001; Laurent, Leung, Maidment, & Balleine, 2012; Bertran-Gonzalez, Laurent, Chieng, Christie, & Balleine, 2013; Laurent, Bertran-Gonzalez, Chieng, & Balleine, 2014; Morse et al., 2020; Laurent & Balleine, 2021). Further, the metabolic function of the NAcC and the cognitive function of the NAcS identified in PIT tasks are disrupted in obesity. In obese-prone rats, action performance in the presence of food-predictive cues is amplified (Derman & Ferrario, 2018; Derman & Ferrario, 2020). In these same rats and in people with obesity, the ability to use food-predictive cues to select actions is impaired (Derman & Ferrario, 2020; Lehner, Balsters, Bürgler, Hare, & Wenderoth, 2017). Interestingly, these abnormal behaviors can be reproduced by manipulating activity of NAc cholinergic interneurons (CINs). Silencing NAcC CINs amplifies general PIT (Collins et al., 2019) whereas genetic ablation of NAcS CINs removes specific PIT (Morse et al., 2020). It has been shown that NAc CINs express InsRs where the binding of insulin increases their excitability (Stouffer et al., 2015). Given its association with reduced InsR sensitivity, brain insulin resistance is likely to decrease the excitability of NAc CINs and could therefore contribute to the metabolic and cognitive deficits displayed by people with obesity in PIT tasks.

The present series of experiments investigated whether alteration of insulin signaling on NAc CINs alters the metabolic regulation of action performance and the cognitive control of action selection in a similar manner as that observed in obesity. We employed two distinct approaches in mice to interfere with insulin signaling on NAc CINs. The first involved a high-fat diet (HFD) model of obesity, which has been shown to broadly reduce InsR expression in several brain regions, including the NAc (Fetterly et al., 2021; Clegg et al., 2011; Sims-Robinson et al., 2016). The second approach was a genetic technique that selectively excised InsRs from cholinergic cells, including NAc CINs. We then examined in these mice the metabolic regulation of food-seeking using a general PIT task or the cognitive control of action selection using a specific PIT task. Post-mortem analyses were conducted to link potential behavioral deficits to changes in insulin signaling or the overall activity in NAc CINs.

## 2. Methods

### 2.1. Subjects

The subjects were adult (at least 8 weeks old) male and female C57BL6/J mice (76 females and 76 males) obtained from the Animal Resources Center (Perth, Western Australia, Australia). There were also adult male and female ChAT-InsR^-/-^ mice (11 females and 9 males) or ChAT-InsR^+/-^ mice (15 females and 9 males). These transgenic mice were generated by crossing mice expressing Cre-recombinase in choline acetyltransferase (ChAT) positive cells (ChAT-Cre mice; #006410; The Jackson Laboratory, Maine, USA) with mice harboring *loxP* sites flanking exon 4 of the InsR gene (#006955; The Jackson Laboratory). After multiple crossings in animals selected according to their genotype, we obtained ChAT-InsR^-/-^ mice that lacked the InsR on cholinergic cells and ChAT-InsR^+/-^ that carried wild-type InsR expression. Efforts were made to allocate an equivalent number of female and male mice to each experimental group, and separates analyses failed to reveal any influence of sex over behavior (Fs<2.36). All mice were group housed (*n* = 2-5) in plastic cages located in a climate-controlled colony room maintained on a 12 h light/dark cycle (lights on between 7:00 A.M. and 7:00 P.M.). They were provided ad libitum food and water until they were food restricted to maintain them at ∼85% of their free-feeding body weight during behavioral protocols. The Animal Care and Ethics Committee at the University of New South Wales approved all procedures, which took place during the light cycle.

### 2.2. Diet treatment

C57BL6/J mice were randomly assigned to receive ad libitum standard laboratory chow or HFD (Specialty Feeds, Western Australia, Australia). The laboratory chow contained 4.6% fat w/w (12% kilocalories from fat) and was chosen as a more nutritionally complete alternative to a modified low-fat diet. The HFD contained 23.5% fat w/w (45% kilocalories from fat) and was composed of lard (20.7 g/110 g) and soya bean oil (2.8 g/100 g). Diets were based on the American Institute of Nutrition Guidelines (AIN93). Body weight and food intake were measured manually throughout the experiments twice per week between 9:00 A.M. and 12:00 P.M. Food intake data were measured/analyzed per cage of four mice.

### 2.3. EchoMRI

EchoMRI was conducted to assess diet-induced changes to body fat levels. This was conducted in the Mark Wainwright Analytical Centre of UNSW Sydney. EchoMRI uses principles of nuclear magnetic resonance to measure fat, lean mass, free water, and total water mass of the animals. It is a fast and non-invasive way to accurately measure body composition to compliment manual body weight data collected during diet manipulations. Mice were gently restrained using cylindrical holders for 90 s without any sedation or anesthesia. EchoMRI was conducted the day before starting the diet treatment (week 0), on the last day of the diet treatment (week 8), and 5 weeks later (week 13), before starting the behavioral procedures.

### 2.4. Drugs

Standard Humulin NPH insulin (Eli Lilly and Co., Melrose Park, New South Wales, Australia) was dissolved in 0.9% saline and injected intraperitoneally (i.p.). Dosing for peripheral insulin challenges prior to transcardial perfusion was 10 U/Kg. Control i.p. injection used 0.9% saline. All peripheral injection volumes were 5 mL/Kg.

### 2.5. Behavioral apparatus

Training and testing took place in MED Associates chambers (Med Associates, Vermont, USA) enclosed in sound- and light-resistant shells. Each chamber was equipped with a pump fitted to a magazine that could deliver a 20% sucrose solution (0.025 mL). Each chamber was also equipped with two pellet dispensers that could deliver either grain or purified food pellets (20 mg, #F0163 and #F0071, Bio-Serv, New Jersey, USA) in the magazine when activated. Two retractable levers could be inserted to the left or right of the magazine that contained an infrared photo beam to detect head entries. The chambers also included a Sonalert to deliver 3 kHz pure tone and a white noise generator (80 dB). A 3 W, 24 V house light provided illumination of the operant chamber. A set of computers running MED Associates proprietary software (Med-PC V) controlled all experimental events and recorded magazine entries and lever presses.

### 2.6. Behavioral procedures

#### 2.6.1. General Pavlovian-Instrumental Transfer

Mice were first placed in the chambers for 60 min in the absence of any event. They were returned to the chambers the next day and received 4 x 2-min presentations of each stimulus (i.e., tone and noise) in a pseudorandom order. These two pre-exposure sessions aimed to reduce neophobia to the chambers and stimuli. The mice then received 8 sessions of Pavlovian conditioning across 8 consecutive days. In each session, a stimulus (S1; tone or noise) was paired with the delivery of a food outcome (O1; grain or purified pellet). Each session included 6 x 2-min presentations of S1 with a variable intertrial interval (ITI) ranging from 3 to 7 min. The outcome was delivered on a random time (RT) 30-s schedule during the stimulus. The identity of the stimulus and outcome was fully counterbalanced across animals. Head entries into the magazine were recorded during the stimulus and during a period of equal length immediately before (pre-S1).

Next, mice received 8 sessions of instrumental conditioning across 8 consecutive days. In each session, a novel food outcome (O2; purified or grain pellet) could be earned by performing a lever press action (A; left or right lever). Each session ran until 30 outcomes had been earned or until 30 min had elapsed. Lever pressing was continuously reinforced for the first 2 days and was then shifted to a variable interval (VI) schedule. The VI schedule increased every two days with VI-15s for days 3 and 4, VI-30s on days 5 and 6, and VI-60s on days 7 and 8. The identity of the lever and outcome was fully counterbalanced. Lever presses were recorded in all sessions. The day after instrumental conditioning, mice received 6 x 2 min presentations of a novel stimulus (S2; noise or tone) in the absence of any other event to generate a neutral stimulus. This was followed the next day by a 20-min instrumental extinction session during which the trained action was available, but no outcome was delivered. The aim was to reduce baseline instrumental lever press rates.

Finally, the mice received a general PIT test that assessed performance on the trained action in the presence or absence of the stimuli. This was conducted under extinction (no outcome delivered) to prevent immediate feedback. The trained lever was made available for 4 minutes to further decrease baseline instrumental lever press rates and then S1 and S2 were separately presented 4 times with a 3-min fixed ITI. The stimuli lasted 2 min and were presented in the following order: tone-noise-noise-tone-noise-tone-tone-noise. Lever presses were recorded during the stimuli and during the 2-min period immediately preceding the stimuli. Half of the C57BL6/J mice that underwent the general PIT protocol were tested hungry whereas the other half were tested sated (standard chow was given ad libitum for 24-48hrs before the test). By contrast, ChAT-InsR^-/-^ mice and ChAT-InsR^+/-^ mice were tested twice: once hungry, and once sated.

#### 2.6.2. Specific Pavlovian-Instrumental Transfer

Mice received the two pre-exposure sessions described above. They then underwent 8 sessions of Pavlovian conditioning across 8 consecutive days. In each session, two stimuli (S1 and S2; tone or noise) were paired with the delivery of two distinct outcomes (O1 and O2; grain pellet or sucrose solution; S1-O1 and S2-O2). Each stimulus was presented 4 times in a pseudorandom order. The length of the stimuli and ITI, the rate of outcome delivery, and the recordings were identical to those described before. The identity of the stimuli and outcomes was fully counterbalanced.

Next, mice received 2 daily sessions of instrumental conditioning across 8 consecutive days. In one session, one lever press action (A1; left or right lever press) earned O1. In the other session, another lever press action (A2; right or left lever press) delivered O2. The order of the session, the identity of the actions and outcomes were fully counterbalanced. Each session lasted until 20 outcomes had been earned or 20 min had elapsed. On the first 3 days, lever presses were constantly reinforced before being shifted to a random ratio (RR) schedule. RR5 was used on days 4 and 5, whereas RR10 was used from days 6 to 8.

In the experiment involving C57BL6/J mice, a 30-min instrumental extinction session was implemented to reduce baseline instrumental responding. The two trained levers were available, and no outcome was delivered.

Finally, the mice received a specific PIT test that assessed performance on the two trained actions in the presence or absence of the stimuli. The test was conducted under extinction to prevent any immediate feedback. The two levers were made available for 9 minutes before presenting each stimulus 4 times. The parameters were identical to those described for the general PIT test.

### 2.7. Histology

#### 2.7.1. In-situ hybridization

In-situ hybridization (RNAscope; Advanced Cell Diagnostics, Hayward, CA) was used to validate the ChAT-InsR^-/-^ transgenic model. Briefly, brains were excised and snap frozen in liquid nitrogen before slicing at 10-20 µm and being mounted directly onto glass slides. Tissue was fixed for 15 min and then dehydrated in 50%, 70%, and 100% ethanol. A hydrophobic barrier was drawn around the tissue samples before a series of hybridization steps were completed with the target ChAT and InsR probes. Probes were hybridized to a cascade of amplification molecules, resulting in binding of dye-labelled probes visible in various fluorescent channels.

#### 2.7.2. Immunofluorescence

Immunofluorescence was conducted for phospho Akt Ser-473 (pAkt) following an insulin challenge to enable quantification of downstream insulin signaling activity. During the challenge, overnight-fasted mice were transcardially perfused 15 minutes after an i.p. injection of either Standard Humulin NPH insulin (*INS*; 10U/kg, 5mL/kg) or saline (*SAL*; 5mL/kg) as a control. Insulin stimulates the pAkt signaling pathway, so disrupted insulin signaling can be indicated by the inability to activate pAkt following insulin administration. This may itself lead to a reduction in InsR expression and has been demonstrated after HFD feeding (Clegg et al., 2011). Immunofluorescence was also conducted for ribosomal protein S6 p-Ser240–244-S6rp (S6rp), an activity marker for CINs (Bertran-Gonzalez, Chieng, Laurent, Valjent, & Balleine, 2012) on brains that were harvested immediately after the general or specific PIT test.

ChAT immunodetection was used to identify the target markers on CINs. One slice containing the target NAc was used from each animal in each set of immunofluorescences. Primary antibodies used were goat anti-ChAT (1:500; Millipore, MA, USA; #AB144P), rabbit anti240-244 phosphorylated ribosomal protein S6 (1:500; Cell Signaling Technology, MA, USA; #2215), and rabbit phospho Akt Ser-473 (1:2000; Cell Signaling Technology; #4060). Secondary antibodies were donkey anti-goat Cy3 (1:800; Jackson Immuno Research, PA; #AB_2307351) and donkey anti-rabbit AF488 (1:500 or 1:1000; Life Technologies, CA, USA; #A-21206). Individual CINs were imaged at 64X to enable quantification of the somatic distribution of pAkt or S6rp.

Immunofluorescence analysis was conducted using the Fiji Image J software (Schindelin et al., 2012). In two channel images, a mask was drawn around the individual CIN, removing the nucleus, and then overlaid onto the target channel. Mean grey value was then measured in the target channel to quantify the fluorescence of the target protein (pAkt or S6rp) in the CIN soma. A total of 1,165 and 892 CINs were analyzed for the pAkt and S6rp quantification, respectively.

### 2.8. Statistical analyses

All data presented met the assumptions of the statistical test used. All hypotheses were established before data collection and are laid out when reporting the results. The analytic plan was pre-specified, and any data-driven analyses are identified and discussed appropriately. Performance during Pavlovian conditioning was analyzed using an elevation ratio of magazine entries. This ratio was obtained by dividing the total number of magazine entries during the relevant stimulus/stimuli by the addition of that number with the total number of magazine entries in the pre-stimulus/pre-stimuli period [i.e., S/(S + pre-S)]. One elevation ratio per animal was calculated on each day. An elevation of 0.5 indicated that the animals entered the magazine as much during the stimulus/stimuli as outside the stimulus/stimuli (i.e., poor learning). By contrast, an elevation ratio above 0.5 showed that the animals entered the magazine more in the presence the stimulus/stimuli than in its/their absence (i.e., good learning). Responding during instrumental conditioning and the PIT tests was analyzed using the number of lever presses per minute (i.e., lever press rate). The differences between groups were analyzed by means of planned orthogonal contrasts. Within-session changes were assessed by planned linear trend analyses. All these procedures and analyses have been described by Hays (Hays, 1963) and were conducted in the PSY software (School of Psychology, The University of New South Wales, Australia). The Type I error rate was controlled at alpha = 0.05 for each contrast tested. If interactions were detected, follow-up simple effects analyses were calculated to determine the source of the interactions.

## 3. Results

### 3.1. HFD interferes with insulin signaling on accumbal cholinergic interneurons

We first aimed to establish that HFD disrupts insulin signaling on NAc CINs. C57BL6/J mice were fed standard chow diet (group CHOW; n = 8 females and 4 males) or HFD (group HFD; n = 4 females and 8 males) for 8 weeks. A preliminary analysis revealed that that body weights were initially similar in both groups (Diet: F<1.43). The data are therefore presented as percentage change from the pre-diet control weight. The CHOW and HFD groups consumed similar calories overall (Figure 1A; Diet: F<1.25), there was no significant increase over time (Week: F<4.5), regardless of groups (Diet x Week: F<3.38). The lack of change in calories was unexpected and was likely due to food intake being recorded per cages of 4 animals. Inspection of Figure 1A does indeed suggest that caloric intake and overall consumption was higher in HFD mice. In line with this suggestion, HFD mice gained more weight during the 8-week period than CHOW mice (Figure 1B; Diet: F(1,22)=19.52, p<.001). There was a gradual and overall increase in body weight over time (Week: F(1,22)=188.59, p<.001), and this increase was larger in HFD mice (Diet x Week: F(1,22)=41.70, p<.001).

**Figure 1.**
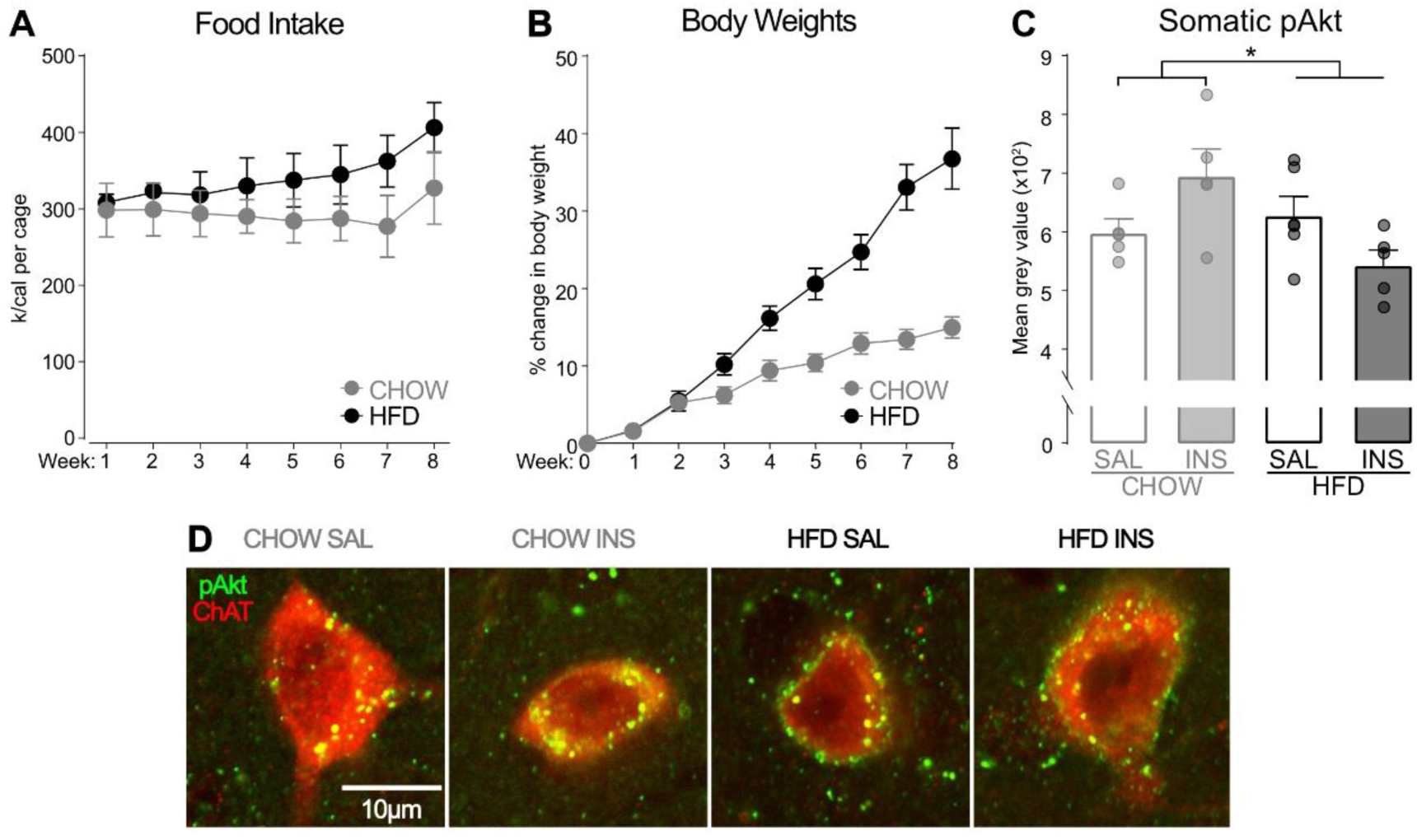
HFD interferes with insulin signaling on accumbal cholinergic interneurons. (A) Cage-averaged food intake was similar across time but (B) HFD mice gained more weight than CHOW mice. (C) NAc CINs from CHOW, but not HFD, mice show insulin-induced pAkt signalling. (D) Representative CINs (red) from the NAc showing pAkt expression (green) for each group of mice. Data are presented as mean +/- SEM. *, p < 0.05.

To determine whether the diet treatment influenced insulin signaling, mice were given an insulin challenge immediately after the end of the treatment. Focusing on the somatic compartment of NAc CINs, we then quantified phosphorylation of the protein Akt (pAkt) (Figure C-D), which is part of the intracellular signaling cascade triggered by InsR activation (Mora, Komander, van Aalten, & Alessi, 2004). Based on previous findings (Arnold et al., 2014; Clegg et al., 2011; Fetterly et al., 2021), we hypothesized that pAkt expression would be disrupted by HFD. Consistent with this hypothesis, there was trend for blunted pAkt activity in HFD mice (Diet: F(1,17)=3.66, p=0.07), indicating reduced insulin function. Although there was no main effect of insulin (Drug: F<1), its impact on pAkt expression depended on the diet treatment (Diet x Drug: F(1,17)=8.01, p<.05). Unfortunately, follow up analyses failed to reveal any significant effect but indicated opposite trends: insulin increased pAkt signaling in the CHOW mice (Drug: F(1,8)=3.78, p=0.09) whereas it decreased this signaling in the HFD mice (Drug: F(1,9)=4.24, p=0.07). Together these results suggest that basal insulin function was reduced in HFD mice, as was the ability for insulin to stimulate its signaling cascade. Both are indicators of insulin resistance. We are therefore confident that HFD blunted insulin signaling in NAc CINs.

### 3.2. HFD produces an abnormally persistent general PIT effect independently of changes in insulin signaling

Having established that a HFD interferes with insulin signaling in NAc CINs, we then used a general PIT task to determine whether HFD consumption influences the capacity of a stimulus predicting food to energize performance of an action earning a different food (Figure 2A). C57BL6/J mice were fed standard chow diet (group CHOW; n = 16 females and 16 males) or HFD (HFD; n = 16 females and 16 males) for eight weeks. The mice were then placed on a chow diet for 5 weeks prior to entering the general PIT task. This was done to produce equivalent training performance in CHOW and HFD mice (Harb & Almeida, 2014). Although we were concerned about potential consequences on HFD-induced changes, InsR downregulation has been shown to persist weeks after HFD removal (Sims-Robinson et al., 2016). CHOW and HFD mice then entered the general PIT task. During Pavlovian conditioning, the mice learned that one stimulus (S1) predicted a particular food outcome (O1) whereas another stimulus (S2) predicted nothing. They then received instrumental conditioning during which one lever press action (A) earned a different food outcome (O2). Finally, a general PIT test assessed the influence of the two stimuli performance of the trained action. Studies employing shifts in primary motivational states (e.g., hunger to satiety) show that metabolic needs modulate general PIT (Corbit et al., 2007; Lingawi et al., 2022; Balleine, 1994; Holland, 2004). To establish whether this modulation was disrupted by the HFD, half of the mice in each diet condition was tested hungry (CHOW-Hungry, n = 8 females and 8 males; HFD-hungry, n = 8 females and 8 males) whereas the other half was tested sated (CHOW-Sated, n = 8 females and 8 males; HFD-Sated, n = 8 females and 8 males).

**Figure 2.**
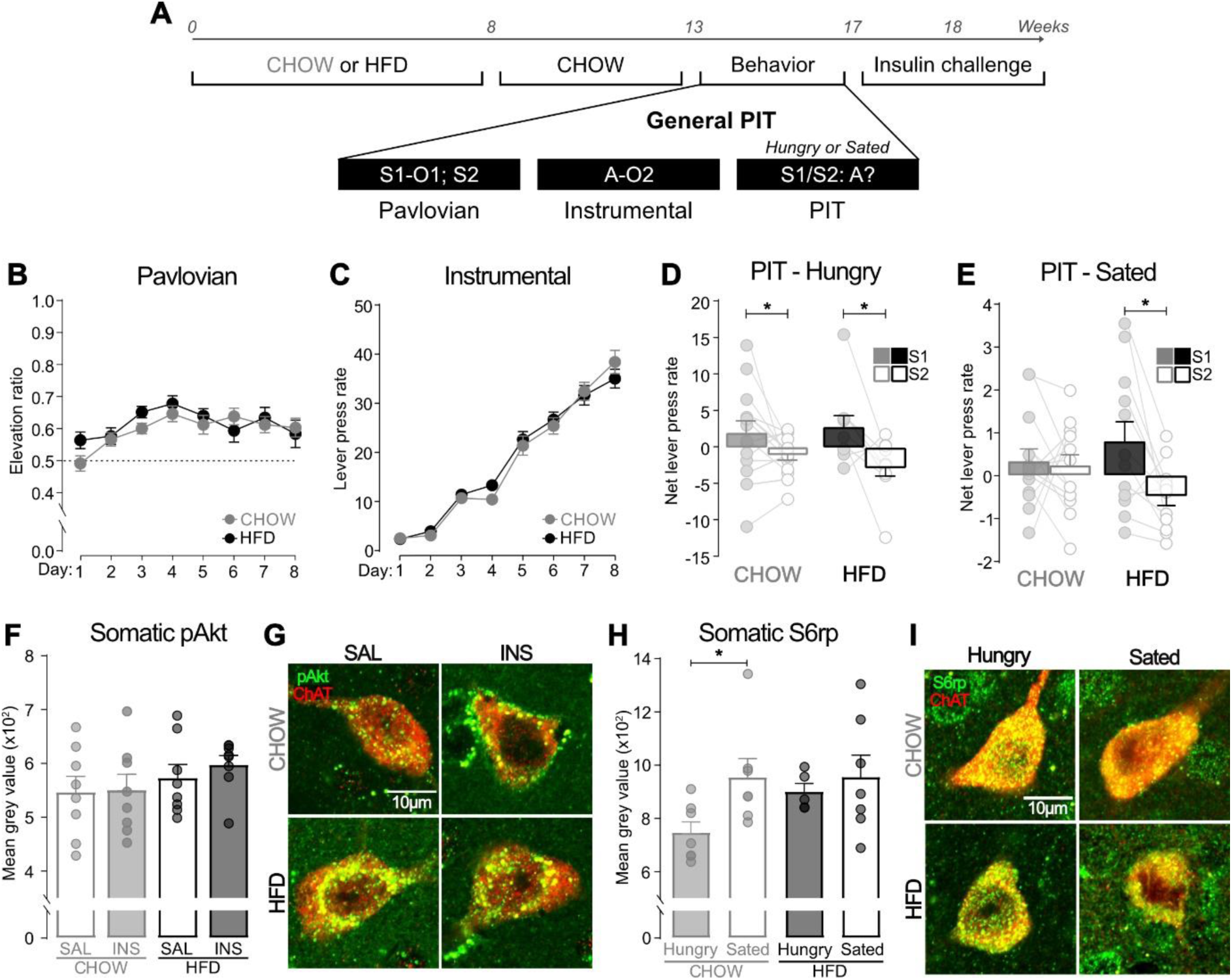
HFD produces an abnormally persistent general PIT effect independently of changes in insulin signaling. (A) Experimental timeline and general PIT design; S1/S2, tone or noise stimuli (counterbalanced); O1/O2, grain and purified pellet outcomes (counterbalanced); A, left or right lever press (counterbalanced). (B) All mice learned that S1 predicted O1. (C) All mice learned to perform a lever press A to earn O1. (D) All hungry mice showed intact general PIT. S1 elevated action performance on A. The neutral stimulus S2 did not influence action performance on response A. (E) When mice were sated, general PIT was abolished in the CHOW group but abnormally preserved in the HFD group. HFD mice showed elevated action performance on A during S1, but not S2, presentations. Neither S1 nor S2 elevated action performance on A for the CHOW group. (F) NAcC CINs showed equivalent pAkt activity across groups and were not responsive to insulin. (G) Representative CINs (red) from the NAcC showing pAkt expression (green) for each group of mice. (H) NAcC CIN activity levels were elevated in the CHOW sated relative to CHOW hungry group. (I) Representative CINs (red) from the NAcC showing S6rp expression (green) for each group of mice. Data are presented as mean +/- SEM. *, p < 0.05.

#### 3.2.1. Behavior

The diet manipulation produced similar changes as in the previous experiment (Supplemental Figure 1A-B) and these were further confirmed by EchoMRI (Supplemental Figure 1C). Separate analyses revealed that the shift in motivational state at test had no impact on data collected prior to test (Fs<2.76). These data were therefore analyzed using only two groups (group CHOW vs. group HFD). Pavlovian conditioning was successful (Figure 2B), and all mice entered the magazine more in the presence of S1 than in its absence, as indicated by an elevation ratio superior to 0.5. This ratio was not influenced by the diet treatment (Diet: F<0.69) and gradually increased across training (Day: F(1,46)=4.06, p<.05), regardless of group (Diet x Day: F<2.03). Instrumental conditioning (Figure 2C) was also successful. Lever press responding was not influenced by the diet (Diet: F<0.11) and it increased as training progressed (Day: F(1,46)=510.17, p<0.001), irrespective of groups (Diet x Day: F<0.51). The training of the neutral S2 and the instrumental extinction session occurred as expected (Supplemental Figure 1D-E).

The data of most importance are those from the general PIT tests (Figure 2D-E), which mice underwent while being hungry (groups CHOW-Hungry and HFD-Hungry) or sated (groups CHOW-Sated and HFD-Sated). It assessed the effects of the trained stimuli (S1 and S2) on performance of the trained action. As expected, satiety severely reduced baseline instrumental responding (i.e., lever press rates in the absence of the stimuli; Satiety: F(1,46)=40.11, p<.001). As such, we analyzed performance during the hungry and sated tests separately. Mice tested hungry displayed similar baseline responding regardless of the initial diet treatment (Diet: F<0.04). Responding during baseline was therefore subtracted from responding during the stimuli to reveal the net effect of the stimuli on performance of the action. Overall net responding (Figure 2D) was not influenced by the diet (Diet: F<0.07), but it was higher in the presence of the food-predictive stimulus S1 than in the presence of neutral stimulus S2 (Stimulus: F(1,22)=12.61, p<0.01), regardless of groups (Stimulus x Diet: F<1.05). Thus, the diet treatment did not disrupt the general PIT effect as the food-predictive stimulus S1 elevated performance of the action that earned another food in all hungry mice. Mice tested sated also displayed similar baseline responding regardless of the initial diet treatment (Diet: F<1.53). We therefore plotted performance as the net effect of the stimuli on instrumental responding again (Figure 2E). Once again, overall net responding was not influenced by the diet (Diet: F<0.13) but, surprisingly it was higher in the presence of the food-predictive stimulus S1 than in the presence of neutral stimulus S2 (Stimulus: F(1,24)=6.52, p<.05). Critically, the difference between the two stimuli depended on the initial diet treatment (Stimulus x Diet interaction: F(1,24)=4.51, p<.05). Simple effect analyses revealed that HFD mice displayed general PIT, as responding in the presence of S1 was higher than in the presence of S2 (Stimulus: F(1,11)=9.33, p<.05). By contrast, this difference was absent in CHOW mice (Stimulus: F<0.11). Taken together, our results confirm that general PIT is sensitive to the test motivational state (Corbit et al., 2007; Lingawi et al., 2022; Balleine, 1994), as it was abolished in sated CHOW mice. However, they also show that HFD removes this sensitivity as general PIT persisted in the HFD group despite the shift to satiety.

#### 3.2.2. Immunofluorescence

Our first experiment demonstrated that HFD treatment interferes with insulin signaling in NAc CINs, and we examined whether a similar disruption could be uncovered in the present experiment. A subset of mice in each experimental group received an insulin challenge followed by pAkt quantification. We focused on CINs the NAcC, as general PIT requires this sub-territory and not the adjacent NAcS (Corbit et al., 2016; Corbit et al., 2001). Our analysis did not distinguish between the motivational states at test (i.e., Hunger vs. Satiety) as there is no reason to believe it would interfere with insulin signaling measured several days later. Unexpectedly, pAkt expression in the somatic compartment of NAcC CINs (Figure 2F-G) was similar in CHOW and HFD mice (Diet: F<2.04) and was not influenced by insulin stimulation (Drug: F<0.31), regardless of groups (Diet x Drug: F<0.16). Thus, unlike our initial experiment, we failed to find any evidence that HFD interfered with insulin signaling on NAcC CINs in mice submitted to the general PIT task.

Our inability to observe changes in insulin on NAcC CINs does not exclude the possibility that activity in these interneurons did modulate the general PIT effect. To test for this possibility, a subset of mice in each experimental group was euthanized immediately after the general PIT test and we quantified S6rp expression in NAcC CINs (Figure 2H-I), which has been found to be a reliable indicator of activity in these interneurons (Bertran-Gonzalez et al., 2012). Expression of S6rp was similar in CHOW and HFD mice (Diet: F<1.42) and this was not influenced by the motivational state of the mice at test (Diet x Satiety: F<1.39). However, there was a trend for greater S6rp expression in mice tested sated relative to mice tested hungry (Satiety: F(1,22)=4.11, p=0.06). We hypothesized that this trend was likely generated by one group failing to display the general PIT effect when tested sated (group CHOW-Sated) while both groups (group CHOW-Hungry and HFD-Hungry) exhibited the effect when tested hungry. This hypothesis is supported by evidence that stimulating NAcC CINs activity abolishes general PIT (Collins et al., 2019). Based on this, we conducted *a priori* simple effect analyses to determine whether the motivational state at test influenced S6rp expression of CHOW and HFD mice differently, given the differences in behavioral performance. Simple effects revealed that satiety at test enhanced S6rp expression in CHOW mice (Satiety: F(1,12)=6.42, p<.05), while it failed to do so in HFD mice (Satiety: F<0.29). Thus, the loss of the general PIT effect in CHOW mice tested sated was linked to an increase in NAcC CIN activity compared to CHOW mice tested hungry. By contrast, the persistence of the effect in HFD mice tested sated was associated with a level NAcC CIN activity that was equivalent to that displayed by hungry HFD mice. Collectively, these results confirmed that NAcC activity opposes general PIT (Collins et al., 2019).

### 3.3. InsR excision from cholinergic cells has no effect on general PIT

The previous experiment demonstrated that HFD treatment freed general PIT from the control exerted by primary motivational states. That is, HFD mice displayed general PIT whether they were tested hungry or sated. This perseverance was linked to changes in NAcC CINs activity, but not to alteration in insulin signaling on the same interneurons. This failure contrasts with our first experiment showing that HFD interfered with NAc CINs response to insulin. Although this discrepancy indicates that some aspect of the dietary reversal procedure employed in the previous experiment rescued insulin signaling in NAcC CINs, it also leaves open the possibility that disrupting this signaling can in fact influence general PIT. To address this possibility, we generated mice that lack InsR expression on cholinergic cells (ChAT-InsR^-/-^ mice; n = 4 females and 3 males) and submitted them to our general PIT task (Figure 3A). We compared performance of these mice with that of their wildtype littermates (ChAT-InsR^+/-^ mice; n = 10 females and 3 males) that harbor normal InsR expression. All mice received two general PIT tests, one while being hungry, and the other while being sated.

**Figure 3.**
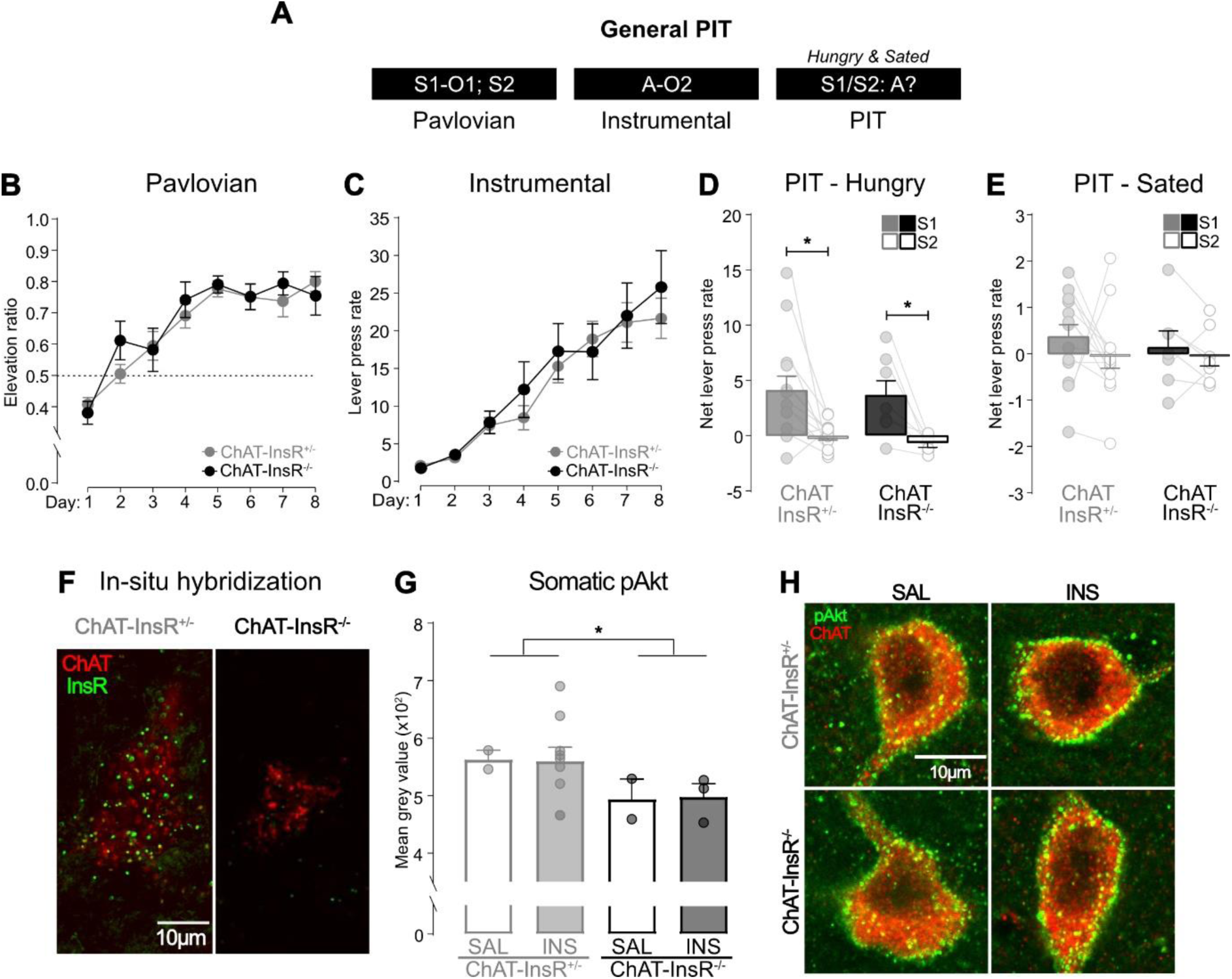
InsR excision from cholinergic cells has no effect on general PIT. (A) Design for general PIT behavioural protocol; S1/S2, tone or noise stimuli (counterbalanced); O1/O2, grain and purified pellet outcomes (counterbalanced); A, left or right lever press (counterbalanced). (B) All mice learned that S1 predicted O1. (C) All mice learned to perform a lever press A to earn O2. (D) All hungry mice showed intact general PIT. S1 elevated action performance on A. The neutral stimulus S2 did not influence action performance on response A. (E) When mice were sated, general PIT was abolished in both groups. (F) In-situ hybridisation confirmed the loss of the InsR in ChAT-InsR^-/-^ mice, and (G) Immunofluorescence indicated that basal pAkt expression was also reduced in ChAT-InsR ^-/-^ mice relative to ChAT-InsR ^+/-^ mice. (H) Representative CINs (red) from the NAcC showing pAkt expression (green) for each group of mice. Data are presented as mean +/- SEM. *, p < 0.05.

#### 3.3.1. Behavior

Pavlovian conditioning was successful (Figure 3B), as the elevation ratio of magazine entries for S1 was superior to 0.5. This ratio was equivalent in both groups (Genotype: F<0.19), it gradually increased across days (Day: F(1,18)=84.25, p<.001), irrespective of groups (Day x Genotype: F<0.19). Instrumental conditioning (Figure 3C) occurred as expected. Lever press responding was similar in both groups (Genotype: F<0.17), and it increased as training progressed (Day: F(1,18)=77.32, p<0.001) regardless of groups (Genotype x Day: F<0.14). The training of the neutral S2 and the instrumental extinction session occurred as expected (Supplemental Figure 2A-B).

The data of most interest are those from the general PIT tests (Figure 3D-E), which mice underwent while being hungry or sated. Because all mice were first tested hungry and then sated, we analyzed the two tests separately. During the hungry test, baseline instrumental responding was similar across groups (Genotype: F<0.23), allowing us to focus on the net effect of the stimuli on instrumental responding. Overall net responding (Figure 3D) was similar in both groups (Genotype: F<0.24) and it was higher in the presence of the food-predictive S1 than in the presence of the neutral S2 (Stimulus: F(1,18)=15.85, p<.001). This difference was present regardless of groups (Stimulus x Genotype: F<0.1), indicating that all mice displayed the general PIT effect. During the sated test, the two groups of mice also displayed similar baseline instrumental responding (Genotype: F<1.48). Overall net responding (Figure 3E) was similar across groups (Genotype: F<0.09). Critically, responding was similar in the presence of S1 and S2 (Stimulus: F<1.09), irrespective of groups (Stimulus x Genotype: F<0.16). Thus, the general PIT effect was abolished in both groups, indicating that motivational states controlled the expression of general PIT whether InsR had been excised from cholinergic cells or not. Together, these results suggest that insulin signaling on NAcC CINs does not modulate the capacity of food-predictive stimuli to enhance performance of an action earning a different food.

#### 3.3.2. In-situ hybridization and immunofluorescence

We employed in-situ hybridization and immunofluorescence to validate the genetic model used in this experiment. In-situ hybridization conducted in a separate cohort of mice (Figure 3F) confirmed the loss of InsRs in NAc CINs in ChAT-InsR^-/-^ mice relative to ChAT-InsR^+/-^ mice. The immunofluorescence analysis involved pAkt expression following an insulin challenge in ChAT-InsR^-/-^ and ChAT-InsR^+/-^ mice submitted to the general PIT tests. It showed that basal pAkt expression was greater in ChAT-InsR^+/-^ mice than ChAT-InsR^-/-^ mice (Figure 3G; Genotype: F(1,12)=7.39, p<.05) but failed to isolate any effect of insulin on pAkt expression. Regardless, the lower basal pAkt signal confirmed InsR excision in NAcC CINs.

### 3.4. HFD has no effect on specific PIT

Having found that HFD consumption regulates the capacity of food-predictive stimuli to energize action performance, we then evaluated whether it also regulates the influence of the same stimuli on action selection in a specific PIT task (Figure 4A). C57BL6/J mice were fed a standard chow diet (group CHOW; n = 16 females and 16 males) or HFD (HFD; n = 16 females and 16 males) for eight weeks. The mice were then placed on a chow diet for 5 weeks prior to entering the specific PIT task. The task started with Pavlovian conditioning during which two stimuli (S1 and S2) predicted two distinct food outcomes (O1 and O2; S1-O1 and S2-O2). The mice then received instrumental conditioning whereby the two outcomes could be earned by performing one of two lever press actions (A1 and A2; A1-O1 and A2-O2). Finally, half of the mice in each diet condition (group CHOW-PIT: n = 8 females and 12 males; group HFD-PIT: n = 12 females and 8 males) was given a specific PIT test that assessed how the stimuli influenced choice between the two actions (S1: A1 vs. A2; S2: A1 vs. A2). The mice were only tested hungry as primary motivational states do not control specific PIT (Corbit et al., 2007; Lingawi et al., 2022; Balleine, 1994; Sommer et al., 2022). The other half of the mice were control mice and their choice between actions was assessed in the absence of the stimuli (group CHOW-CTL: n = 8 females and 4 males; group HFD-CTL: n = 4 females and 8 males).

**Figure 4.**
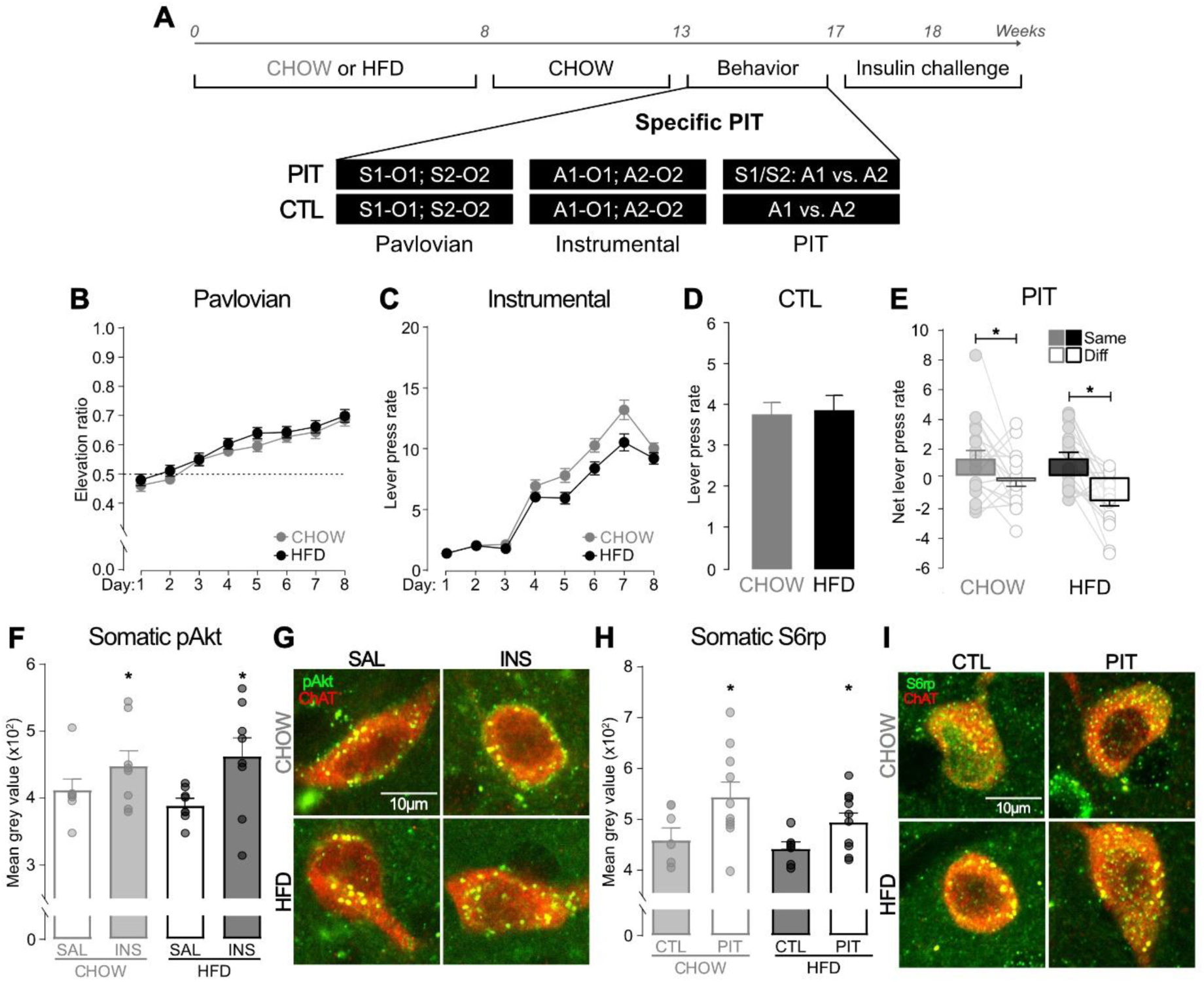
HFD has no effect on specific PIT. (A) Experimental timeline and specific PIT design; S1/S2, tone or noise stimuli (counterbalanced); O1/O2, grain pellet and sucrose solution outcomes (counterbalanced); A1/A2, left and right lever presses (counterbalanced). (B) All mice learned that S1 and S2 predicted O1 and O2, respectively. (C) All mice learned to perform two lever presses, A1 and A2, to earn O1 and O2, respectively. (D) All CTL mice showed similar overall lever press rates on A1 and A2 during the test session. (E) CHOW and HFD mice showed intact specific PIT. S1 biased action selection for A1 and S2 biased action selection for A2. Responding in the absence of the stimuli did not differ between groups. (F) NAcS CINs were responsive to insulin stimulation, with increased pAkt in insulin-versus saline-injected mice. However, this response was not disrupted by the HFD treatment. (G) Representative CINs (red) from the NAcS showing pAkt expression (green) for each group of mice. (H) NAcS CIN activity levels were elevated in PIT groups relative to CTL groups, but activity was not disrupted by the HFD treatment. (I) Representative CINs (red) from the NAcS showing S6rp expression (green) for each group of mice. Data are presented as mean +/- SEM. *, p < 0.05.

#### 3.4.1. Behavior

The data concerning the impact of the diet confirmed previous observations (Supplemental Figure 3A-C). Separate analyses revealed that the presence or absence of stimuli at test had no impact on data collected prior to this test (Fs<1.25). These data were therefore analyzed using only two groups (group CHOW vs. group HFD). Pavlovian conditioning was successful (Figure 4B), as indicated by an elevation ratio superior to 0.5. The ratio was equivalent in both groups (Diet: F<1.24), and it increased across days (Day: F(1,56)=142.38, p<.001), irrespective of groups (Diet x Day: F<0.08). Instrumental conditioning was also successful (Figure 4C), as responding on the two actions increased across days (Day: F(1,56)=645.23, p<.001). However, this increase was larger in CHOW mice than in HFD mice (Diet x Day interaction: F(1,56)=4.79, p<.05) while overall responding followed the same pattern (Diet: F(1,56)=4.80, p<.05). Yet, responding did increase as training progressed in both groups (CHOW-Day: F(1,27)=396.92, p<.001; HFD-Day: F(1,29)=360.36, p<.001), indicating that all mice learnt that the two actions earned the two outcomes. All mice received an instrumental extinction session before test and this session occurred without incident (Supplemental Figure 3D).

The data from the test are presented in Figure 4D-E. We first consider the control groups of mice (groups CHOW-CTL and HFD-CTL) that were presented with the two levers without the two stimuli. As can be seen in Figure 4D, overall rate of lever pressing during the entire test session was similar regardless of the diet (Diet: F<0.04). The data of most interest are those from the two groups of mice (groups CHOW-PIT and HFD-PIT) that received the specific PIT test. Baseline instrumental responding was similar in both groups (F<0.001), allowing us to analyze the net effect of the stimuli on instrumental performance (Figure 4E). This net effect is reported when the stimulus predicted the same outcome as the action (Same) and when it predicted the different outcome (Different; Diff). Overall responding was similar regardless of the diet treatment (Diet: F<1.70) and it was higher on the same action than on the different action (Stimulus: F(1,36)=24.22, p<.001). Critically, the diet did not influence the selection of the same action relative to the different action (Stimulus x Diet interaction: F<2.77), indicating that all groups displayed the specific PIT effect. Thus, all mice preferentially selected the action earning the same outcome as that predicted by the stimulus.

#### 3.4.2. Immunofluorescence

As before, we implemented an insulin challenge in a subset of mice and quantified pAkt expression (Figure 4F-G). We focused our quantification on CINs located in the NAcS, as they are the ones to be required for specific PIT (Morse et al., 2020). Somatic pAkt expression was similar in CHOW and HFD mice (Diet: F<0.03) and it was increased by insulin administration (Drug: F(1,27)=6.43, p<.05), regardless of groups (Diet x Drug: F<0.56). Thus, we failed to find evidence that the HFD interfered with insulin signaling on NAcS.

In another set mice, we analyzed S6rp expression in the somatic compartment of NAcS CINs immediately after behavioral test (Figure 4H-I). In this analysis, the critical comparison was between the groups that showed specific PIT (groups CHOW-PIT and HFD-PIT) and those that could not (groups CHOW-CTL and HFD-CTL). As expected, S6rp expression was higher in mice displaying specific PIT than in mice that did not (Test, PIT vs. CTL: F(1,27)=7.13, p<.05), regardless of whether these mice received the HFD treatment or not (Test x Diet: F<0.41). Overall, the diet had no impact on S6rp expression (Diet: F<1.67). These findings are consistent with the view that NAcS CINs are required for specific PIT.

### 3.5. InsR excision from cholinergic cells has no effect on specific PIT

The previous experiment revealed that the HFD treatment failed to interfere with specific PIT and insulin signaling on NAcS CINs, as measured by pAkt expression. It left opened the question as to whether insulin functioning on NAcS CINs modulates specific PIT. To address this question, the present experiment submitted ChAT-InsR^-/-^ and ChAT-InsR^+/-^ to specific PIT (Figure 5A). The protocol was identical to that described before, except that the control groups that were only tested with the trained actions were omitted.

**Figure 5.**
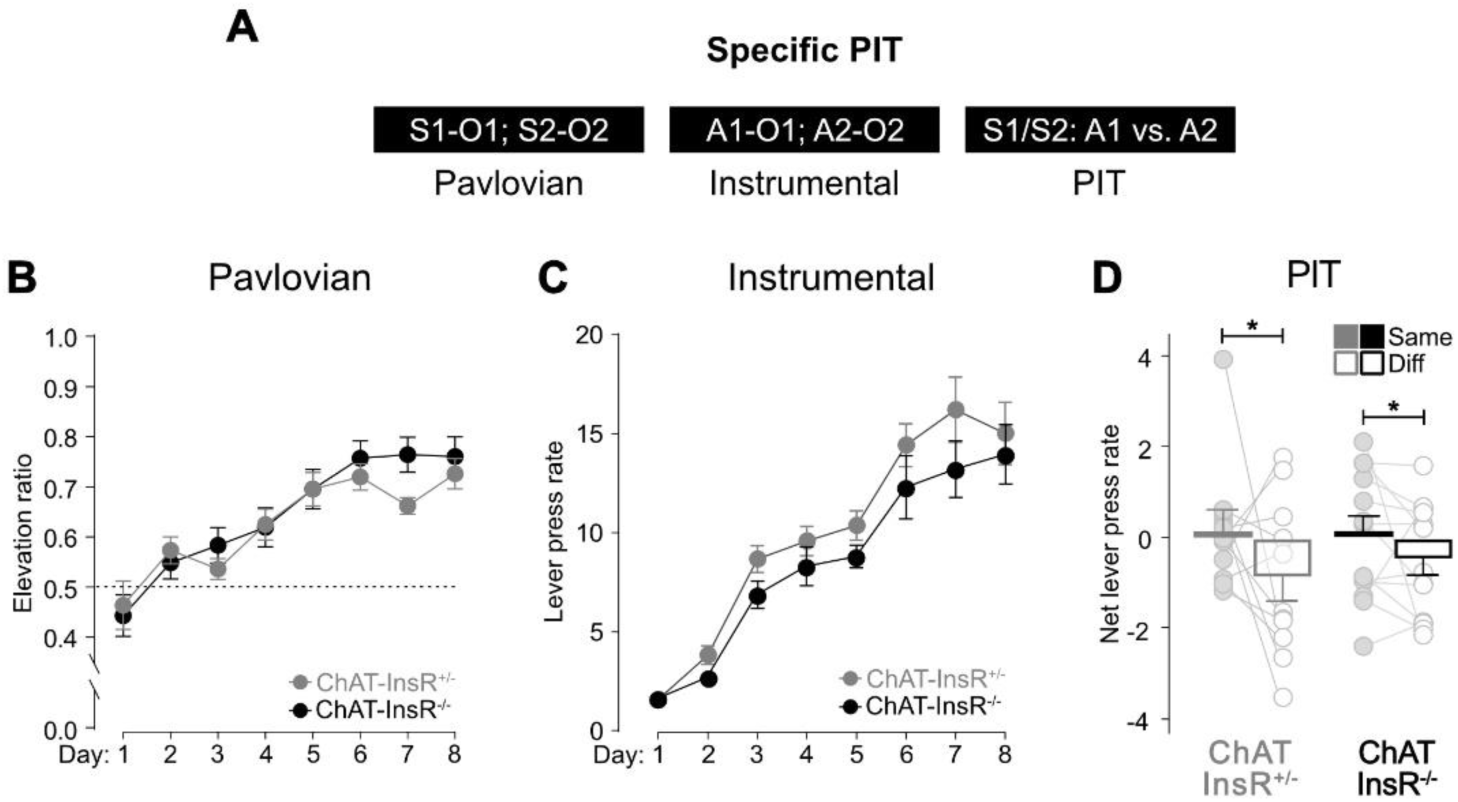
InsR excision from cholinergic cells has no effect on specific PIT. (A) Design for specific PIT behavioural protocol; S1/S2, tone or noise stimuli (counterbalanced); O1/O2, grain pellet and sucrose solution outcomes (counterbalanced); A1/A2, left and right lever presses (counterbalanced). (B) All mice learned that S1 and S2 predicted O1 and O2, respectively. (C) All mice learned to perform two lever presses, A1 and A2, to earn O1 and O2, respectively. (D) Both groups showed intact specific PIT. S1 biased action selection for A1 and CS2 biased action selection for A2. Data are presented as mean +/- SEM. *, p < 0.05.

Pavlovian conditioning occurred smoothly (Figure 5B). The elevation ratio of magazine entries was superior to 0.5, indicating that the mice entered the magazine more in the presence of the stimuli than in their absence. There was no overall effect of genotype (Genotype: F<0.42), the elevation ratio gradually increased across training (Day: F(1,22)=113.09, p<.001) and this increase was irrespective of groups (Genotype x Day: F<2.36). Instrumental conditioning (Figure 5C) was successful and overall lever press responding increased as training progressed (Day: F(1,22)=142.62, p<.001). This increase was not influenced by genotype (Genotype x Day: F<0.41), which did not modulate overall performance (Genotype: F<2.10).

The data of most interest are those from the specific PIT test (Figure 5D). Baseline instrumental responding was similar across groups (Genotype: F<0.73). The data are therefore plotted as net responding on the same or different action. Overall net responding was similar across groups (Genotype: F<0.08) and it was higher on the same action than on the different action (Stimulus: F(1,22)=4.44, p<.05). This was true regardless of genotype (Stimulus x Genotype: F<0.53), indicating that all mice preferentially selected the action earning the same outcome as that predicted by the stimulus. Thus, InsR excision from cholinergic cells spared the specific PIT effect, suggesting that this effect does not require insulin-mediated modulation of NAcS CINs.

## 4. Discussion

The present experiments examined whether insulin signaling on NAc CINs regulates the capacity of food-predictive stimuli to 1) energize action performance in a general PIT task, and 2) guide action selection in a specific PIT task. Two strategies were implemented to interfere with insulin function: a HFD treatment and InsR excision from cholinergic cells. We found that the general PIT effect was left intact by HFD when mice were tested hungry. However, HFD produced a persistent form of the effect that survived a shift from hunger to satiety. The persistence of general PIT was associated with NAcC CINs activity but was not linked to local disruption of insulin function. Accordingly, InsR excision from cholinergic cells had no effect on general PIT whether it was assessed under hunger or satiety. Next, we found that the HFD had no impact on specific PIT and did not interfere with insulin signaling in NAcS CINs. However, successful specific PIT was associated with sustained activity in NAcS CINs. Finally, InsR excision on cholinergic cells spared specific PIT, confirming that insulin function on NAcS CINs is unlikely to contribute to this effect.

General PIT was left intact by HFD in mice tested hungry. In this condition, the food-predictive stimulus energized performance of the instrumental action that had been trained to deliver another food, whether the mice had received the diet treatment or not. This lack of HFD effect reveals important differences between preclinical models of obesity. Indeed, previous research has shown that obese-prone rats displayed an amplified general PIT effect (Derman & Ferrario, 2018; Derman & Ferrario, 2020). This amplification has been proposed to originate from changes in glutamatergic or insulin function on spiny-projection neurons in the NAc (Derman & Ferrario, 2018; Fetterly et al., 2021). This suggests that our HFD treatment probably failed to generate these physiological alterations, thereby leaving general PIT unaffected when the mice were tested hungry. Our treatment did however have an impact on general PIT. When the mice were tested sated, HFD mice continued to display general PIT whereas control mice did not. This result indicates that HFD overrides the traditional control exerted by primary motivational states over the expression of general PIT (Corbit et al., 2007; Lingawi et al., 2022; Balleine, 1994). This strengthens the validity of HFD consumption as a preclinical model of obesity, as this disorder is characterized by overeating whereby food is consumed even in the absence of metabolic needs (Ferrario, 2020; Vainik, García-García, & Dagher, 2019; Johnson, 2013).

The perseverance of general PIT produced by HFD under satiety was not associated with alterations in insulin signaling. Insulin-induced pAkt expression in NAcC CINs was not affected by the HFD treatment in mice submitted to the general PIT task. This contrasts with our initial observation that the same treatment interfered with insulin function on NAc CINs, when assessed immediately upon cessation of the diet. There are important methodological differences that are likely to be responsible for these discrepant findings. First, the general PIT experiment included a 5-week diet reversal whereby mice were placed on a standard chow diet after the 8-week HFD treatment. The reversal was implemented to guarantee equivalent performance during the training stages of the general PIT task, as evidence indicates that HFD can reduce such performance (Harb & Almeida, 2014). This reduction would have complicated any interpretation of potential differences during the general PIT test. Importantly, there was reason to believe that the 5-week diet reversal would have minimal impact, as InsR downregulation has been shown to persist at least 8 weeks after HFD removal (Sims-Robinson et al., 2016). A second difference that may account for the discrepant findings is the addition of the food restriction schedule necessary to study general PIT. Food restriction has been shown to enhance striatal sensitivity to insulin (Stouffer et al., 2015). Regardless of whether one or both differences did contribute, sensitivity to insulin in NAcC CINs was rescued in HFD mice submitted to general PIT, implying that local insulin signaling did not promote the capacity of HFD to free general PIT from the control exerted by primary motivational states. This was further confirmed by our genetic model, as mice lacking InsR on cholinergic cells exhibited general PIT when tested hungry but not when tested sated. This experiment convincingly demonstrates that general PIT is not controlled by insulin signaling on NAcC CINs, underscoring the need for future experiments to investigate the neurobiological mechanisms underlying the perseverance of general PIT following HFD consumption. Although insulin-related changes in NAcC CIN function do not appear to be involved in general PIT, we found evidence that activity in these interneurons plays a critical role in regulating this effect. This evidence was gathered by quantifying S6rp expression in the somatic compartment of NAcC CINs, which has been shown to be a reliable marker of their activity (Bertran-Gonzalez et al., 2012). We found that the loss of general PIT in control mice tested sated was linked to an increase in the S6rp signal in NAcC CINs relative to control mice tested hungry. By contrast, the S6rp signal was similar in HFD mice whether they were tested hungry or sated. Given that these mice displayed general PIT in both conditions, whereas control mice only exhibited general PIT when tested hungry, we reason that NAc CINs activity hinders the expression of general PIT. This is entirely consistent with a recent study demonstrating that stimulation of NAc CINs abolishes, whereas their silencing enhances, general PIT (Collins et al., 2019).

Specific PIT was left intact by the HFD treatment. In CHOW control and HFD-treated mice, the stimuli guided choice towards the action with which they shared a common food outcome. The lack of HFD effect once again reveals important differences between preclinical models of obesity. Indeed, specific PIT has been shown to be impaired in junk-food fed or obesity-prone rats (Kosheleff et al., 2018; Derman & Ferrario, 2020). The source of the difference remains unclear but could again be attributed to the incapacity of our HFD treatment to produce the physiological changes triggered by junk-food consumption or displayed by obesity-prone rats. It is worth noting, however, that the human literature has also been inconclusive with respect to the relationship between specific PIT and obesity, with some finding impairment, enhancement or no alteration at all (Lehner et al., 2017; Meemken & Horstmann, 2019; Watson, Wiers, Hommel, & de Wit, 2014; Watson, Wiers, Hommel, & de Wit, 2014). Clearly, additional studies are required to clarify under which condition specific PIT can be observed, impaired, or enhanced in obesity and the neurobiological changes controlling these effects. Nevertheless, the present study indicates that our HFD treatment left intact the cognitive control of action selection modelled by specific PIT. Consistent with the general PIT experiment, mice given the HFD and submitted to specific PIT did not exhibit a distorted response to insulin on NAcS CINs. However, it is noteworthy that NAcS CINs showed increased pAkt expression following insulin administration, whereas the same interneurons in the NAcC did not. This is consistent with the view that insulin has distinct and potentially competing effects within the mesolimbic circuitry and NAc sub-territories (Ferrario & Reagan, 2018), and that the rate of transport of insulin into the brain varies across brain regions (Rhea, Rask-Madsen, & Banks, 2018). As noted, our incapacity to observe disruption of insulin signaling on NAcS CINs was likely due to the 5-week diet reversal implemented after the HFD and/or the food restriction required to study specific PIT. Regardless, our last experiment revealed that InsR excision from cholinergic cells had no impact on specific PIT. We are therefore confident that insulin signaling on NAcS CINs is not required for the capacity of food-predictive stimuli to guide action selection. Yet, we obtained convincing evidence that NAcS CIN activity is necessary for this process. Somatic expression of S6rp in NAcS CINs was elevated in mice displaying specific PIT relative to control mice that could not show the effect. This result is in line with previous studies (Bertran-Gonzalez et al., 2013; Laurent et al., 2014; Morse et al., 2020) and confirms that NAcS CINs are required for specific PIT.

In summary, the present experiments demonstrate that the capacity of food-predictive stimuli to energize action performance and to guide action selection does not recruit insulin signaling on NAcC CINs and NAcS CINs, respectively. They do however reveal that the energizing effect of the stimuli is regulated by NAcC CINs activity whereas their guiding effect is modulated by NAcS CINs activity. Nevertheless, the main finding rests in the observation that a HFD treatment overrides the typical control exerted by primary motivational state over the expression of general PIT. That is, the treatment allowed the food-predictive stimuli to energize the performance of action earning another food even in the absence of hunger. This mimics a key characteristic of obesity whereby food is consumed in the absence of metabolic needs, confirming the validity of HFD as a tool to model some of the symptoms associated with this disorder. Future studies are needed to isolate the neural changes that allows HFD to distort the typical control exerted by food-predictive stimuli over action performance.

## 5. Author contributions

J.M.G., N.W.L., B.L., M.D.K., B.C.C. and V.L. conceived and designed the experiments. J.M.G. conducted the experiments with assistance from N.W.L and B.L. Data were analysed and interpreted by J.M.G. and V.L. The manuscript was written by J.M.G. and V.L.

