## Supplemental Figures for "High fat diet allows food-predictive stimuli to energize action performance in the absence of hunger, without distorting insulin signaling on accumbal cholinergic interneurons"

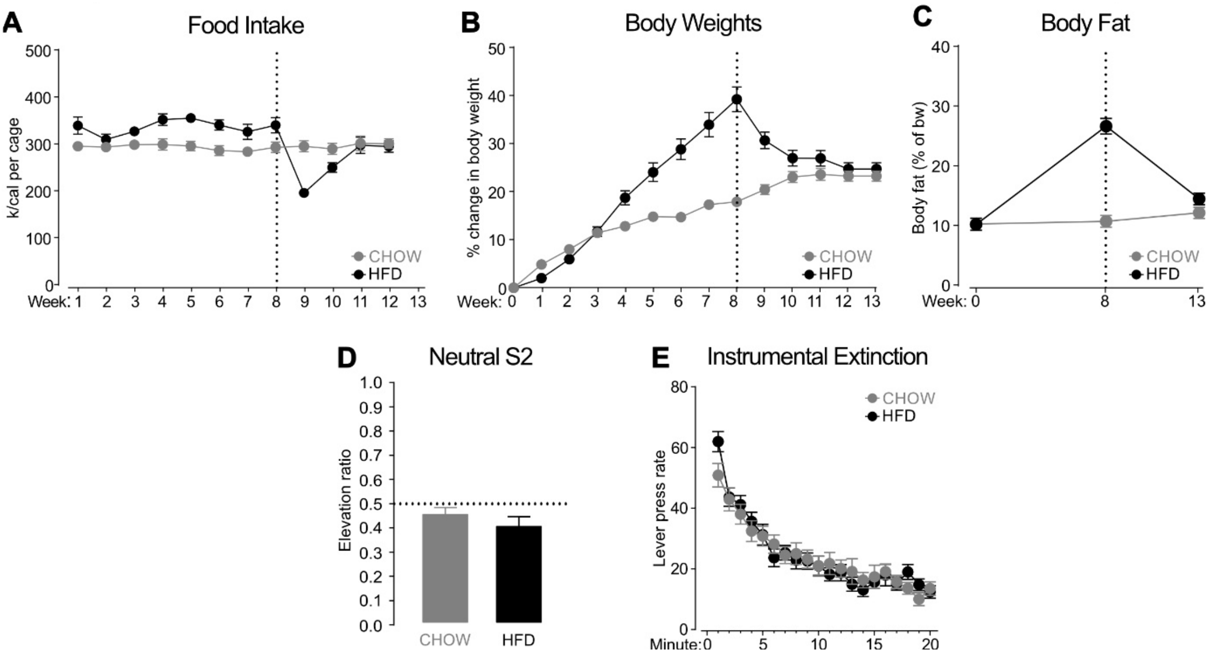


**Figure S1. Related to Figure 2. Food intake, body weight, and additional training data.** Mice were either given a chow or high fat diet for the first 8 weeks (weeks 1-8). All mice then received the chow diet for 5 weeks (weeks 9-13) before starting the food deprivation schedule for behavioural training. Week 13 food intake data were removed from statistical analyses due to only being recorded for part of the week. Analyses revealed that the shift in motivational state at test had no influence on food intake, body weights, or body fat at any stage of the diet treatment (Satiety: Fs<2.76). Thus, the data were analyzed using two groups (group CHOW vs. group HFD). At week 0 body weights were similar in CHOW and HFD mice (Diet; F<0.01), so they are thus presented in B) as percentage change in body weight from the pre-diet control weight. **(A-B)** In the first 8 weeks, the HFD mice consumed more calories than the CHOW mice (Diet: F(1,11)=9.23, p<.05). The number of calories consumed remained stable over time (Week: F<0.26) but was greater in the HFD mice (Diet x Week interaction: F(1,11)=5.07, p<.05). During this 8-week period, there was a steady increase in body weights (Week: F(1,60)=266.15, p<.001) across all mice (Diet: F<3.60), but this increase was larger in the HFD mice than the CHOW mice (Diet x Week interaction: F(1,60)=41.80, p<.001). All mice were then placed on the chow diet for a period of 5 weeks before food deprivation. The CHOW mice consumed more calories than the HFD mice (Diet: F(1,12)=4.44, p<.05), and there was an overall increase in calories consumed (Week: F(1,12)=53.13, p<.001). However, HFD mice showed a greater change in calories consumed over these final weeks (Diet x Week interaction: F(1,12)=38.06, p<.001). The severe drop in food intake in the HFD mice seen in week 9, upon the dietary reversal to chow, is not unusual. HFD fed mice have been shown to exhibit diminished preference for and less consumption of chow food, relative to chow fed controls, but this response is not permanent (Altherr et al., 2021). During the 5-week period there were overall changes in body weights (Week: F(1,60)=6.94, p<.05), with greater gross changes in the HFD mice (Diet: F(1,60)=5.22, p<.05). As was expected, this pattern of body weight changes was different in the HFD versus CHOW mice (Diet x Week interaction: F(1,60)=39.54, p<.001). These changes were evident as a decrease in body weights for the HFD mice (Week: F(1,30)=24.70, p<.001) due to the dietary reversal to the less calorie dense chow, and a continued age-related increase for the CHOW mice (Week: F(1,30)=17.19, p<.001). **(C)** Prior to starting the diet treatment body fat was similar in both groups (Diet; Fs<0.02). Over the three timepoints, body fat was higher in HFD mice than in CHOW mice (Diet: F(1,60)=75.83, p<.001) and body fat increased across time (Week: F(1,60)=45.07, p<.001). However, this increase was not linear and depended on whether mice had been given the chow or HFD (Diet x Week interaction: F(1,60)=7.04, p<0.01). There was an increase in body fat over time in both the CHOW mice (Week: F(1,30)=18.48, p<.001) and HFD mice (Week: F(1,30)=28.23, p<.001), but the magnitude of these increases is greater in the HFD mice. At the end of the diet treatment, HFD mice had higher body fat than CHOW mice (Diet: F(1,60)=111.67, p<.001). This difference persisted after all mice had been placed on the chow diet (Diet: $F(1,60)=8.53$, $p<0.01$), even though the size of the difference was severely reduced. **(D)** All mice treated S2 as a neutral stimulus. The overall number of magazine entries was similar in all groups (Diet, Satiety; $Fs<1.53$). Entries did not differ according to the presence or absence of S2, as indicated by an elevation ratio inferior to 0.5, irrespective of groups (Diet, Satiety; $Fs<1.01$). **(E)** Instrumental extinction was successful and lever press responding on the trained action declined as training progressed (Minute: $F(1,46)=249.93$, $p<.001$). This decline was similar across groups (Minute x Diet, Minute x Satiety; $Fs<1.10$) as was overall responding (Diet, Satiety; $Fs<2.41$).

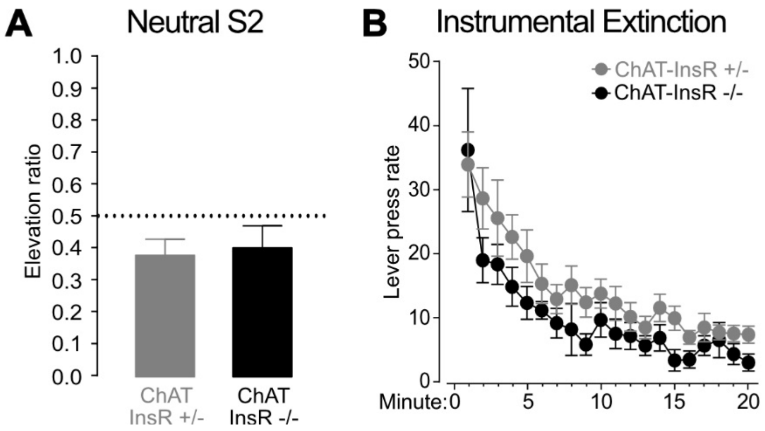


**Figure S2. Related to Figure 3. Additional training data.**

**(A)** All mice treated S2 as a neutral stimulus. The overall number of magazine entries was similar in both groups (Genotype: $F<0.86$). Entries did not differ according to the presence or absence of S2, as indicated by an elevation ratio inferior to 0.5, irrespective of group (Genotype: $F<0.08$). **(B)** Instrumental extinction was successful and lever press responding on the trained action declined as training progressed (Minute: $F(1,17)=31.12$, $p<.001$). This decline was similar across groups (Minute x Genotype: $F<0.12$), as was overall responding (Genotype: $F<2.29$).

### Supplemental Figure 3. Related to Figure 4. Food intake, body weight, and additional training data.

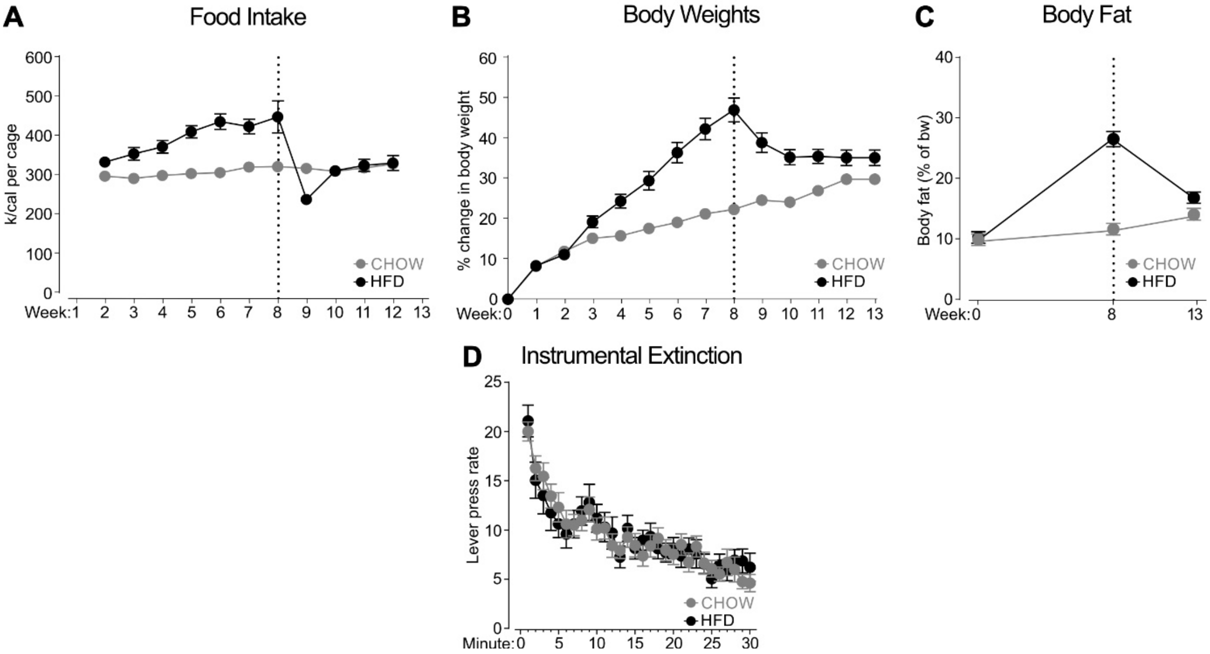


**Figure S3. Related to Figure 4. Food intake, body weight, and additional training data.**

Week 1 and 13 food intake data were omitted from analyses due to experimenter error and incomplete data. Analyses revealed that the future test received (CTL or PIT) did not influence food intake or body fat (Test; $Fs<3.54$) at any stage of the diet manipulation, however it did interact with baseline body weights. At week 0 before starting the diet treatment, body weights were similar in CHOW and HFD mice (Diet: $F<0.08$), and CTL and PIT mice (Test; $F<0.19$), but overall the groups were not equal (Test x Diet interaction: $F(1,60)=4.57$, $p<.05$). Separate analyses revealed no differences between CHOW and HFD mice at week 0 (Diet, Diet x Test; $Fs<0.36$), instead that the interaction was due to sex imbalances between the groups. A greater proportion of males led to the CHOW PIT (Sex: $F(1,36)=105.29$, $p<.001$) and HFD CTL (Sex: $F(1,20)=128.67$, $p<.001$) groups heavier at week 0. Thus, the changes observed as a result of the diet treatment are presented as CHOW groups versus HFD groups. Body weight data for the 13-week diet treatment are presented as percentage change in body weight from the pre-diet control weight. **(A-B)** In the first 8 weeks, the HFD mice consumed more calories than the CHOW mice (Diet: $F(1,11)=46.68$, $p<.001$). The number of calories consumed increased over time (Week: $F(1,11)=7.78$, $p<.05$) in all groups (Diet x Week; $F<2.39$). During this 8-week period, there was a steady increase in body weights (Week: $F(1,60)=342.78$, $p<.001$), but the HFD mice gained more weight (Diet: $F(1,60)=28.25$, $p<.001$) and at a faster rate (Diet x Week interaction: $F(1,60)=61.46$, $p<.001$) than the CHOW mice. All mice were then placed on the chow diet for a period of 5 weeks before food deprivation. Over these final weeks, HFD and CHOW mice consumed the same amount of calories (Diet: $F<0.40$), and there was a gradual increase in calories consumed (Week: $F(1,11)=42.24$, $p<.001$), however the pattern of consumption varied in order to reach the same total calories (Diet x Week interaction: $F(1,11)=25.01$, $p<.001$). The severe drop in food intake in the HFD mice seen in week 9 upon the dietary reversal to chow is standard in these protocols and was observed and discussed previously. There was an initial decrease in food intake in the HFD mice, which stabilises with time resulting in the same total calories consumed between groups. During the 5-week period there were overall changes in body weights (Week: $F(1,60)=8.50$, $p<.01$), with greater gross changes in the HFD mice (Diet: $F(1,60)=32.028$, $p<.001$). Unsurprisingly, this pattern of body weight changes was different in the HFD versus CHOW mice (Diet x Week interaction: $F(1,60)=101.09$, $p<.001$). These changes were evident as a decrease in body weights for the HFD mice (Week:

F(1,30)=48.14, p<.001) as a result of the dietary reversal to the less calorie dense chow, and a continued age-related increase for the CHOW mice (Week: F(1,30)=100.81, p<.001). **(C)** Prior to starting the diet treatment, body fat was similar in both groups (Diet: Fs<0.08). Over the three timepoints, body fat was higher in HFD mice than in CHOW mice (Diet: F(1,60)=41.05, p<.001) and body fat increased across time (Week: F(1,60)=58.20, p<.001). The difference in this increase over time between the groups was only marginally significant (Diet x Week interaction: F<3.25, p=0.08). Inspection of the figure clearly indicates that there was a gradual increase in body fat over time in the CHOW groups compared to a severe increase in the HFD groups over the first 8 weeks, followed by a drop in body fat in the HFD groups in the last 5 weeks. **(D)** Instrumental extinction was successful and lever press responding on the trained actions A1 and A2 declined as training progressed (Minute: F(1,56)=138.35, p<.001). This decline over time was similar across both diet groups (Minute x Diet: F<0.01) and overall responding was similar across diet groups too (Diet: Fs<0.20). Despite identical press rates in CTL and PIT groups during instrumental conditioning, CTL groups pressed more during extinction than PIT groups (Test: F(1,56)=4.38, p<.05), but this did not depend on the diet (Diet x Test interaction: F<1.21).
